# Assessment of exposure to Di (2-ethylhexyl) phthalate (DEHP) metabolites and Bisphenol A (BPA) and its importance for the prevention of cardiometabolic diseases

**DOI:** 10.1101/2021.12.06.470607

**Authors:** Fabrizia Carli, Demetrio Ciociaro, Amalia Gastaldelli

## Abstract

Exposomics analyses have highlighted the importance of biomonitoring of human exposure to pollutants, even non-persistent, for the prevention of non-communicable diseases like obesity, diabetes, non-alcoholic fatty liver disease, atherosclerosis and cardiovascular diseases. Phthalates and bisphenol A (BPA) are endocrine disrupting chemicals (EDCs) widely used in industry and in a large range of daily life products that increase the risk of endocrine and cardiometabolic diseases especially if the exposure starts during childhood. Thus, it is important the biomonitoring of exposure to these compounds not only in adulthood but also in childhood. This was the goal of the LIFE-PERSUADED project that measured the exposure to phthalates (DEHP metabolites, MEHP, MEHHP, MEOHP) and BPA in Italian mother-children couples of different ages. In this paper we describe the method that was set up for the LIFE PERSUADED project and validated during in the proficiency test (ICI/EQUAS) showing that accurate determination of urinary phthalates and BPA can be achieved starting from small sample size (0.5ml) using two MS techniques applied in cascade on the same deconjugated matrix.

## 1. Introduction

Plasticizers are colorless and odorless esters, mainly phthalates or bisphenols, that increase the elasticity of a material (e.g., polyvinylchloride (PVC). For these properties they are widely used in industry and in a large range of daily life products. The majority of these are considered as non-persistent pollutants and have a short half-life. However, the exposure to these agents has significant consequences on human health since once ingested or inhaled are harmful for health since they act as endocrine disrupting chemicals (EDCs), i.e., they are able to interfere with the endocrine system by modulating and/or disrupting the metabolic and hormonal functions responsible for the maintenance of homeostasis, reproduction, development and/or behavior [1]. As many studies have shown, they increase the risk to develop endocrine and cardiometabolic diseases such as obesity [2, 3], non-alcoholic fatty liver disease (NAFLD) [4–7], diabetes [8–11], beta cell function [12–14], hypertension [15–18], atherosclerosis [13, 19, 20], coronary artery disease (CAD) [21–24], chronic kidney disease (CKD) [25–27], thyroid dysfunction [28–30] (Figure 1). The longer the exposure (e.g., prenatal or during childhood) the higher the risk [31].

**Figure 1.**
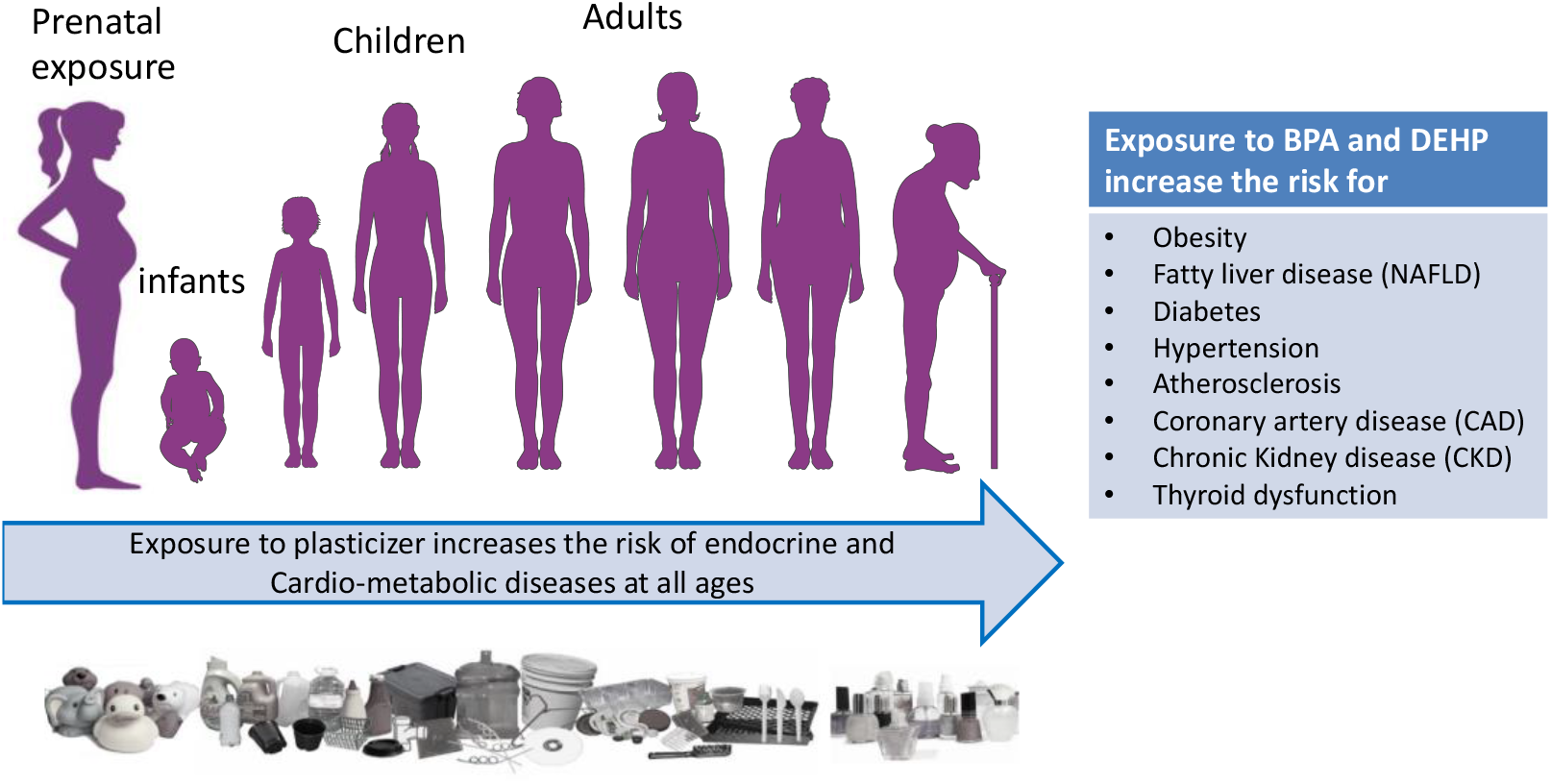
Long term exposure to phthalates and BPA increases the risk of cardiometabolic diseases.

### 1.1. Fate of phthalates and bisphenols in humans

Phthalates are derived from phthalic acid and are used primarily to make plastic products more flexible. For the past 50 years, phthalate production has increased and plastic products may contain up to 40-50% of phthalate by weight. Phthalates are present in all consumer products containing plastics including packaging materials for food, children’s and baby’s toys, household items, waterworks and paints but also in medical devices, such as tubing and intravenous bags and in personal product care such as cosmetics and perfumes [32, 33]. Phthalates are bound not covalently to the polymer matrix making them highly susceptible to leaching [34] so they can easily migrate from plastic to air and food, but also to blood and skin. That explains why human exposure to phthalates and bisphenols is so widespread and nearly ubiquitous, with more than 75% of the U.S. population had phthalate metabolite detected in urines (more recent references) [35, 36]. Once absorbed into the human body phthalates are rapidly metabolized with half-lives within few hours [34, 36]. Their metabolism depends on the structure formula i.e., the short-er-chain-length phthalates undergo preferentially to a phase-I metabolism in which they are hydrolyzed to the corresponding monoesters, then conjugated with glucuronic acid in a phase II metabolism and excreted into urine and feces; the longer-chain-length phthalates after first hydrolyzed and then oxidized. After absorption DEHP is rapidly metabolized by hydrolysis to monoester, mono(2-ethylhexyl) phthalate (MEHP) [37] that is further metabolized to mono(2-ethyl-5-hydroxyhexyl) phthalate (MEHHP) and mono(2-ethyl-5-oxohexyl) phthalate (MEOHP) (Figure 2). A second step is hepatic glucuronic conjugation that is important to reduce their potential biological activity as well as to facilitate their excretion from the human body increasing their water solubility [36, 38–40]. The great majority of DEHP metabolites is found in urine in glucuronidated form.

**Figure 2.**
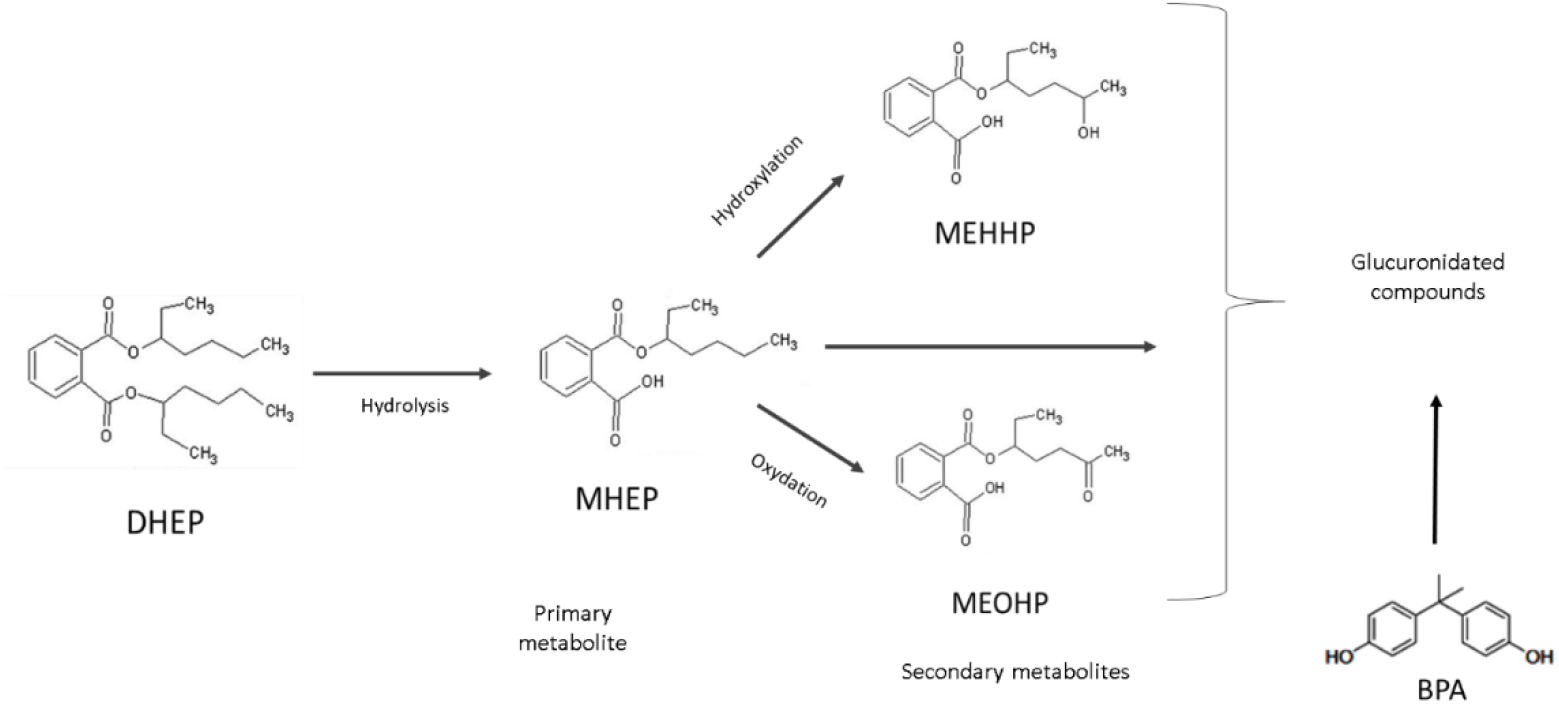
Metabolism of DEHP and BPA. DEHP is hydrolyzed in MEHP and is again hydrolyzed in MEHHP or is oxidated in MEOHP; the three metabolites are conjugated with glucuronic acid. BPA is directly conjugated with glucuronic acid.

Another group of plasticizers are bisphenols that include Bisphenol A (BPA), and the new Bisphenol S and F. Bisphenols contain two hydroxyphenyl functionalities, most of them based on diphenylmethane (Figure 2). BPA is a monomeric building block of polycarbonate and is widely used as an additive to plastics (for example polyvinyl chloride) and other consumer products, such as thermal paper [34]. Thus, food and drink packaging are an important source of BPA that can migrate to the food because some monomers are unbound and factors such as elevated temperature favors leaching process [41]. After absorption, Bisphenol A is mostly glucuronidated in liver and then excreted in urines [42, 43] (Figure 2). Thus, total (i.e., free plus glucuronidated) urinary BPA and phthalate metabolites MEHP, MEHHP, MEOHP are considered the biomarkers for assessing human exposure to BPA and to parent phthalate DEHP [37].

### 1.2. Disrupting action of phthalates and bisphenols

Phthalates and bisphenols are among the most studied endocrine disrupting chemicals (EDCs). This disruption can happen through the alteration of normal hormone levels, halting or stimulating the production of hormones, or changing the way hormones travel through the body, thus affecting hormones control. Moreover, they can modify the individual epigenetic characteristics through DNA methylation, histone modification and microRNAs causing alteration in gene expression with adverse health effects [44]. Moreover, several studies suggest that phthalates and BPA contribute to the increase in obesity [3, 45], development of insulin resistance and type 2 diabetes [46–48] and renal dysfunction as recently our results have highlighted founding a positive association between exposure to DEHP and degree of albuminuria in subjects with type 2 diabetes [49]. Phthalates can bind and activate several nuclear receptors, particularly in the liver and adipose tissue but also in other organs as thyroid and in reproductive system [50, 51]. Phthalates, as many EDCs, act as antagonists or agonists of nuclear hormone receptors (NRs). Phthalates are selective modulators of peroxisome proliferator-activated receptors (PPARs), a family of nuclear receptors involved in glucose and lipid metabolism. In humans three PPAR subtypes have been identified: PPARα and PPARβ/δ mainly involved in the fatty acid catabolism in muscle and liver; PPARγ involved mainly in lipid metabolism and adipogenesis. [52, 53]. Activation of PPARα in the liver by DEHP and MEHP modifies protein expression leading to an increase in cell proliferation [54]. In the presence of ligands, the PPARs form a heterodimer with the retinoid X receptor (RXR) modulating the expression of target genes presenting PPAR-response elements in the promoter region for adaptation to the metabolic state. The monoesters of phthalates, such as the MEHP derived from DEHP, have been indicated also as activators of PPARγ [54–56]. The phthalate monoester MEHP binds PPARγ similar to Rosiglitazone, an antidiabetic drug in the thiazolidinedione (TZD) class, but this binding induces the recruitment only a subset of coregulators activating subset of target genes with adipogenic effect [53, 54]. Recent studies have shown that DEHP is able to interfere with the insulin signal by reducing the molecules involved in its signal transduction such as IRS1 and Akt and with hepatic glucose uptake and transcriptional regulation of GLUT2 suggesting DEHP exposure impairs insulin signal transduction and alters glucoregulatory events leading to the development of type 2 diabetes in F1 male offspring [57, 58]. Phthalates and DEHP cause toxicity in many organs: for example, in liver they alter enzyme activities or gene expression and induce oxidative stress [59, 60], induce hepatomegaly and hepatocarcinogenesis [61]; in brain PPARγ hyperactivation leads to apoptosis in undifferentiated neurons [62]. Thus, total daily intake (TDI) for DEHP has been set to 50 μg per kilogram body weight per day (EFSA 2005) while EPA has set the Reference Dose (RfD) to 20 μg per kilogram body weight per day (mg/kg/d) based on increased relative liver weights in guinea pigs [63]. BPA is also able to bind to PPARs. Activation of PPARs by BPA induces insulin resistance in all organs, glucose intolerance, hepatic hypertriglyceridemia, adipogenesis and excess release of non-esterified fatty acids, similarly to high fat diet [64]. BPA exposure can also induce epigenetic modifications associated to type 2 diabetes and obesity. In Sprague–Dawley rats BPA induced the down-regulation of glucokinase (gck) gene in liver leading to glucose intolerance and insulin resistance [65, 66]. BPA was shown to induce epigenetic modifications in genes involved in lipid metabolism: subcutaneous injection of BPA in mice decreased the WAT (white adipose tissue) mRNA expression of Srebpc1, PPARα and Cpt1β and the hepatic expression of CD36 [64]; oral exposure of BPA in male C57BL/6 mice reduced expression of miR-192 with consequent upregulation of Srebf1 and other genes of lipid synthesis, that increase de novo lipogenesis and display NAFLD-like phenotype [67]. Moreover, when pregnant Sprague–Dawley rats were exposure on BPA, in sperm of male F1 and islets of male F2, Igf2 was hypermethylated, decreased its expression and in male F2 offspring there was beta-cell dysfunction [68] DEHP and BPA are also able to activate estrogen receptor ERα and Erβ that are involved in growth and development but also in numerous physiological functions [69, 70]. BPA compared to the natural ligand 17β-estradiol (E2), BPA binds ER receptors with less affinity [71] with the consequences depending on the target tissue, as the effect of interference on cell death and proliferation equilibrium by Erα/ Erβ signaling [72]. Recent data also indicate that BPA is able to bind to another estrogen receptor, 7-transmembrane estrogen receptor, GPR30 expressed in different cell types and cancer cell lines [51]. This receptor is overexpressed in breast and ovarian cancer and GPR knockout female mice are protected from obesity, blood glucose intolerance, and insulin resistance induced by high fat diet [73]. Low doses of BPA are able to increase GPR30 and adipose tissue inflammation with the increase of IL8, IL6, and MCP-1α [51].

### 1.3. Biomonitoring studies

The biomonitoring studies aim at the measurement of the exposure to pollutants in the general population by measuring environmental chemicals, or their metabolites, in biological samples. Environmental chemicals are normally present in the matrix at trace levels, so the first step is to define which biological matrix is the most appropriate, e.g., urine, blood, saliva etc. Urines are more easily collected than blood, e.g., for children, and are preferred for non-persistent environment exposure as BPA and phthalates since their concentration is more abundant in urines where they are excreted as both free and glucuronidated compounds compared to other matrices [74]. Exposure is assessed by the measurement of both free and glucuronic compounds. Typically, exposure is measured in a single spot urine sample (first void urine), but to reduce variability it would be more appropriate 24h urine collection or pool of different samples [75]. The analysis of presence of environmental chemicals is usually done by mass spectrometry, either liquid chromatography mass spectrometry (LC/MS) or gas chromatography mass spectrometry. The concentration of the environmental chemicals is usually done by comparing the peak area of the chemical with the peak area of the corresponding labelled internal standard added at known concentration to the matrix. When spot urines are used for the analysis, the chemical concentrations are normalized to the urinary concentration of creatinine. The preparation and purification of the sample and the choice chromatographic approach have an impact on the quality of the measurement.

## 2. Results

### 2.1. Measurement of DEHP metabolites by LC/MS

The analysis of DEHP metabolites was carried out using a UHPLC/QTOF as described above

#### 2.1.1. Recovery and matrix effect of DEHP metabolites (Table 1)

The recovery of MEHP, MEOHP and MEHHP ranged between 74-97% for both unlabelled and labelled standards. Background levels in blank, measured by adding only internal labelled standards, was 0.28 ng/ml for MEHP and no detectable (ND) for the other metabolites. Matrix effect (ME%) for DEHP metabolites in LC/MS analysis was 90% for MEHP, 102% for MEHOP and 105% for MEHHP.

**Table 1.** Recovery of DEHP metabolites and BPA in blank and urine samples.

|  | MEHP | MEOHP | MEHHP | BPA |
| --- | --- | --- | --- | --- |
| Spike (ng/ml) | 3.8 | 3.8 | 3.8 | 10.4 |
| Blank (ng/ml) | 0.28±0.01 | ND | ND | 0.69±0.006 |
| Urine+spike (ng/ml) | 4.16±0.15 | 3.75±0.20 | 4.60±0.13 | 9.6±0.17 |
| Recovery % | 94% | 79% | 102% | 91.99% |
| Matrix Effect % | 90% | 102% | 105% | NA |

#### 2.1.2. Limit of Detection (LOD) and Quantification (LOQ) of DEHP metabolites (Table 2)

Limit of Detection (LOD) and Quantification (LOQ) were determined for MEHP, MEHHP and MEOHP. LOD for DEHP metabolites ranged between 0.11 to 0.28 ng/ml and LOQ from 0.24 to 0.58 ng/ml. Method Detection Limit (MDL) was the lowest for MEOHP (0.096 ng/ml) and the highest for MEHP (0.238 ng/ml).

**Table 2.** Limit of Detection (LOD), Quantification (LOQ) and Method Detection Limit (MDL); SNR: signal-to-noise ratio

| DHEP metabolites | LOD (ng/ml) | LOQ (ng/ml) | MDL (ng/ml) | SNR |
| --- | --- | --- | --- | --- |
| MEHP | 0.28 | 0.58 | 0.238 | 11.30 |
| MEOHP | 0.18 | 0.48 | 0.096 | 14.50 |
| MEHHP | 0.11 | 0.24 | 0.131 | 10.60 |

#### 2.1.3. Working range and Linearity of DEHP method

For each compound (either MEHP, MEOHP or MEHHP) we tested the linearity and stability of the instrumental measurement by running labelled standard curves once per week before starting the analyses. Each curve contained 5 points where the ^13^C labelled internal standard was at the same concentration used in urine samples while the unlabelled standard was added at increasing concentrations (4.1, 6.1, 8.1, 16.2, 50.7 ng/ml). The standard curves showed good linearity (R>0.995 for MEHP, MEOHP and MEHHP) over the range 4-50 ng/ml (Figure 3).

**Figure 3.**
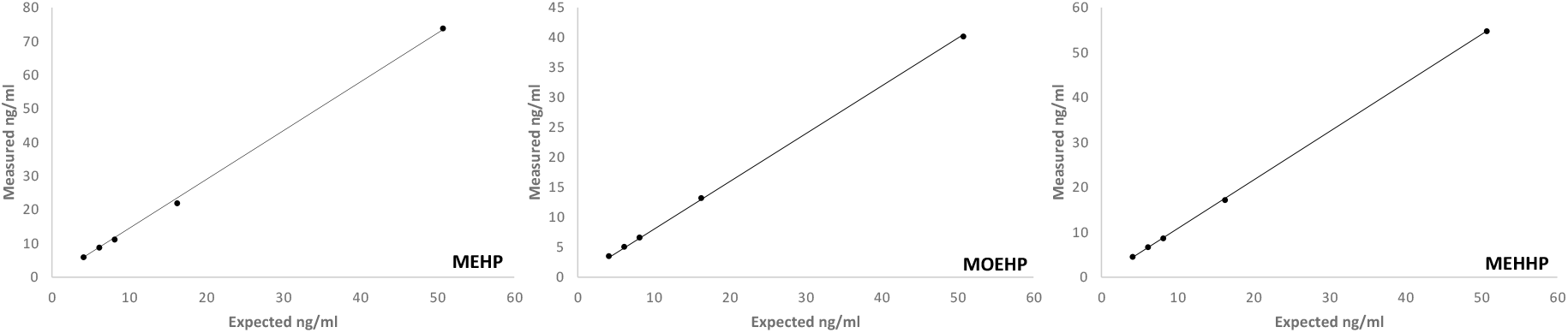
Standard curves of DEHP metabolites in LC/MS. Measured concentrations expressed as ng/ml (n=29) of DEHP metabolites are reported in the y axis while expected (theoretical) values are located in the x axis; R=0.99 for all metabolites.

#### 2.1.4. Stability, reproducibility of the analysis and precision of DEHP method

Several aliquots of quality control (QC) samples were prepared from a urine pool as describe in the methods and analyzed during every run. Figure 4 reports the concentrations of MEHP, MEOHP and MEHHP in QC samples measured for the LIFE-PERSUADED analyses from 2015 to 2018. Concentrations of DEHP metabolites in QC samples showed stability and reproducibility over time as showed by the three years’ plot in Figure 4. MEHHP variability was the highest, though less than 10%. The Relative Standard Deviation (RSD) of DEHP metabolites in QC samples was 6.49% for MEHP, 3.69% for MEOHP and 8.21% for MEHHP.

**Figure 4.**
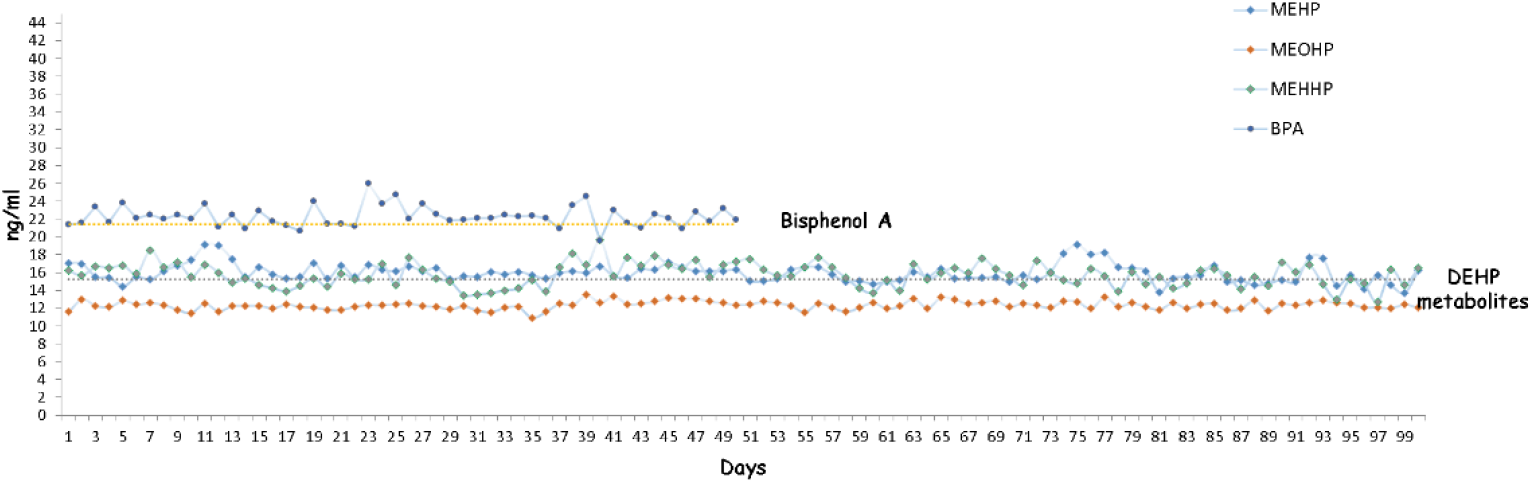
Reproducibility of the matrix QC spiked with MEHP, MEHHP and MEOHP and measured by LC/QTOF from 2015 to 2018 (n=100). The average values ± standard deviation (ng/ml) measured for quality control (QC) samples of standard metabolites were: MEHP: 15.96 ± 1.04; MEHHP: 15.73 ± 1.31; MEOHP: 12.34 ± 0.45 respectively. For BPA we assessed stability and reproducibility in n=50 QC together with DEHP metabolites in LC/MS analysis and average value was 22.30 ± 1.17 ng/ml and RSD=5.25%.

### 2.2. Measurement of Bisphenol A (BPA) by UHPLC/QTOF and GC/MS

Total BPA in urine samples was quantified first using by high resolution UHPLC/QTOF, i.e., during the same run of DEHP metabolites, and then by the method approved by HBM4EU Project (https://www.hbm4eu.eu/online-library/?mdocs-cat=mdocs-cat-20&mdocs-att=null#), that uses GC/MS.

#### 2.2.1. Working range and Linearity of BPA methods

The linearity of both LC/MS and GC/MS methods was tested by preparing a standard curve covering the range from 1.1 to 71.53 ng. The standard curves showed good linearity in both instruments, R = 0.99 (Figure 5).

**Figure 5.**
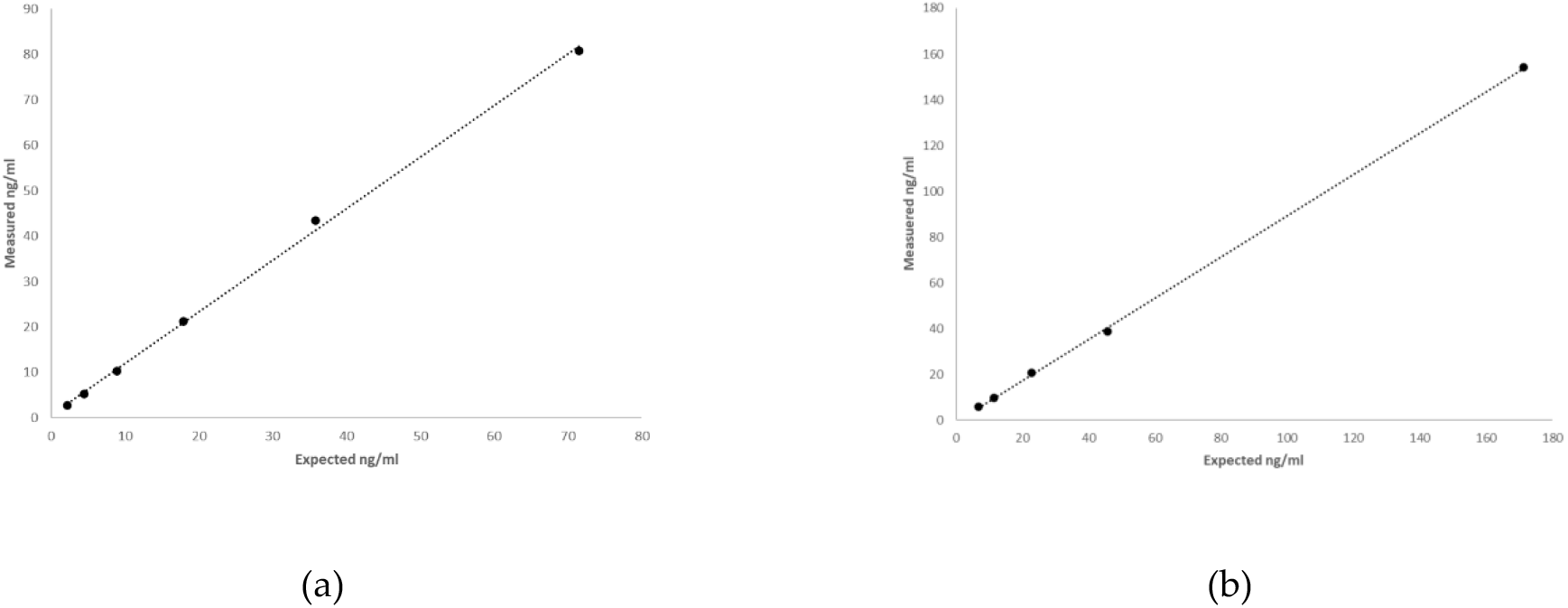
Standard curves of BPA in GC/MS (A) and in LC/MS (B). Measured concentrations expressed as ng/ml were reported in the ordinate axis while expected values were located in the abscissa axis and R=0.99 for both analyses.

#### 2.2.1. Stability, reproducibility of the analysis and precision of BPA methods

Quantification of QC samples for BPA showed good reproducibility, recovery and accuracy by both LC/MS and GC/MS analyses. QCs were measured regularly during GC/MS run and showed good reproducibility and accuracy. QCs (n=100) had a mean of 24±4.3 ng/ml with RSD% of 17.96%. Recovery of d16BPA, added in the same amount in free and deconjugated urine, was much lower in deconjugated urine than in free urine; furthermore, the addition of increasing concentrations of IS after deconjugation did not result in a proportional increase in the signal which, on the contrary, remained very low indicating strong signal suppression. Moreover, the LC/MS analysis of the deconjugated samples showed that the peak area of d16BPA was variable and low, a) despite being added in constant concentrations and b) compared to the IS areas of the DEHP metabolites that also had been added to the same matrix in constant concentrations and analyzed under the same instrumental conditions. Since derivatization of the same samples (deconjugated and non-deconjugated) with GC/MS analysis did not detect the same problems, we concluded that it is likely that compounds created during deconjugation affect sample recovery and ionization.

#### 2.2.2. Limit of Detection (LOD) and Quantification (LOQ) of BPA by GC/MS

We calculated LOD and LOQ according to U.S.EPA procedure (U.S. EPA 2017) and were 0.157 ng/ml and 0.523 ng/ml, respectively. For BPA contamination we analyzed a sample of d16BPA to quantify the blank signal. To stabilize the background signal of the BPA we made 20 successive injections of the internal standard until reaching the minimum blank signal corresponding to 0.69 ng/ ml.

### 2.3. Validation of methods and application in the HUMAN BIOMONITORING STUDY

The analytical methods for DEHP metabolites and BPA were validated during the proficiency test (ICI/EQUAS), organized within the activities of the HBM4EU project, which provided urine samples with reference values from certified laboratories. The analyses were conducted twice in two different analytical batches (about 5 months apart) in double preparation and in double run and the analysis was repeated after about another 5 months. The participants did not know they were two reputable batches. We thus obtained 9 replicates, for low concentration and high concentration sample, which include all possible sources of variability. Relative Standard Deviation (RSD%) ranging from 7.55 until a maximum of 23.91 in a low concentration of MEOHP. MEHP was under limit of quantification a low concentration sample. Trueness of measure was reported as BIAS% and ranging from 5.14 to 12.96 (Table 3).

**Table 3.** Quantification of DEHP metabolites and BPA in 2 reference samples with low and high concentration of metabolites; 9 replicates were measured for DEHP metabolites and 8 for BPA and the average was reported. RDS%= relative standard deviation; LOQ= limit of quantification.

|  | Concentration | Ref Value<br>ng/ml | Replicates | Average<br>ng/ml | Precision<br>(RSD%) | Trueness<br>(Bias%) |
| --- | --- | --- | --- | --- | --- | --- |
| MEHP | Low | 0.567 | 9 | <LOQ | - | - |
|  | High | 5.89 | 9 | 6.65 | 14.43 | 12.96 |
| MEOHP | Low | 1.74 | 9 | 1.63 | 23.91 | 6.22 |
|  | High | 14.6 | 9 | 13.32 | 7.55 | 8.74 |
| MEHHP | Low | 4.12 | 9 | 3.71 | 14.96 | 9.87 |
|  | High | 32.3 | 9 | 30.64 | 13.01 | 5.14 |
| BPA | Low | 0.763 | 8 | 0.751 | 19.17 | 1.55 |
|  | High | 8.4 | 8 | 6.456 | 13.29 | 23.14 |

We were then invited to participate to the ring study organized by the HBM4EU Project together with other 24 laboratories from 16 countries to harmonize bisphenols analysis. Our analytical method was tested during the proficiency test (ICI/EQUAS), during four rounds of intercalibration laboratory tests. We repeated analysis of a low concentration sample and a high concentration sample to assessed RSD% and BIAS% of our analysis for BPA and results are show in Table 3.

## 3. Discussion

The European Commission (EFSA) and the US Environmental Protection Agency (EPA) [76–80] have warned and restricted the use of Di(2-ethylhexyl) phthalate (DEHP) in food packaging and personal care products but they are still widely present in many consumer products [33, 81]. Moreover, in December 2019 EPA designated DEHP as a High-Priority Substance and the chemical is currently undergoing risk evaluation. The creation of materials suitable for the needs of consumer products led to the synthesis of new chemicals that unfortunately are dangerous for human health since they interfere with the hormonal system [82]. In 1997 the Environmental Protection Agency (EPA) de-fined the term endocrine disrupting chemicals (EDC) as “an exogenous agent that interferes with the production, release, transport, metabolism, binding, action, or elimination of natural hormones in the body responsible for the maintenance of homeostasis and the regulation of developmental processes” (EPA 1997). Since then, more and more reports showed how these EDCs act in organisms and the dangers they entail for human health. Great attention has been given to exposure to these substances in growing children where EDCs can act easily and lead to developmental abnormalities. The biomonitoring of these substances allows to understand the exposure to a given pollutant and lead to a regulation of these substances to limit damage to health and increase national health expenditure. Despite restrictions in the employment of DEHP and BPA and replacement by other plasticizers (e.g., dioctyl terephthalate (DEHT), bisphenol S and F), DEHP and BPA are still widely present in packaging and personal care products [33] and they can also easily pass to food, as shown in a study that analyzed food samples taken from fast food restaurants finding DEHP and BPA in more than 70% samples [81]. Although mass spectrometry is the preferred method for the detection of environmental pollutants, the type of chromatographic method used may vary (i.e., gas or liquid chromatography). Moreover, there are several types of mass spectrometers, for LC/MS triple quadrupole LC/MS/MS that acquires data in multiple reaction monitoring (MRM) [83–89], but also hybrid triple quadrupole linear ion trap mass spectrometer (QTRAP) in selected reaction monitoring (SRM) mode [90], providing a good sensitive response. Several studies used GC/MS, with single quadrupole or tandem mass spectrometer, that requires derivatization of the sample, but recently also eliminating the derivatization step [91–93]. At the time of the start of the LIFE PERSUADED project there was no published method using high resolution LC/QTOF. High resolution mass spectrometry is the best method for discovery of known and unknown compounds in the matrix of interest; given its selectivity and accuracy it is becoming a useful tool to study the exposure to EDCs even in small biological sample [94]. Unlike LC/MS/MS system with target MRM acquisitions, the LC/QTOF system allows to acquire and quantify several compounds with high resolution in a single analysis compatibly with the chromatographic method. The method developed for the European project LIFE-PERSUADED project [95] and here reported used LC/QTOF for the evaluation of DEHP metabolites and GC/MS for BPA in urine samples. For both methods there was good agreement with published. The methods were tested and were validated during the proficiency test (ICI/EQUAS), organized within the activities of the HBM4EU project, which provided urine samples with reference values from certified laboratories. The initial setup of the method included the simultaneous measurement of DEHP metabolites and BPA by UHPLC/QTOF, as also recently proposed [96]. We found that the method gave reliable responses for free BPA, but we detected significant signal suppression due to matrix effects after deconjugation for the measurement of total environmental chemicals (i.e., free plus glucuronidated compounds). The analyses were then repeated by GC/MS where the matrix effect was absent. Since the LIFE-PERSUADED study (“Phthalates and bisphenol A biomonitoring in Italian mother-child pairs: link between exposure and juvenile diseases”) involved children, there was a need to start from small urine volumes. Although urine samples are easy to collect in adults, in children this is not always so simple, especially when they are very young; thus, the use of a very sensitive instrument allows the detection of these compounds starting from low volumes. Thus, the method was setup in samples as low as 0.5ml. Our method has the advantage to be easy to perform and shows high stability and reproducibility of the quality control samples measured over 3 years. The method has also been tested and validated during the proficiency test in Interlaboratory Comparison Investigations and External Quality Assurance Schemes (ICI/EQUAS) of the HBM4EU Project showing good agreement with reference values (https://www.hbm4eu.eu/online-library/?mdocs-cat=mdocs-cat-20&mdocs-att=null#). The quality of the measurement of environmental chemicals in biological matrices is determinant for the biomonitoring studies but often challenging. Measurements of phthalates or bisphenols concentrations in biological matrices are not currently standardized. In 2018 HBM4EU organized an Inter Laboratory Comparison Investigation (ICI) to compare data and methods within several laboratories (18 for phthalates and 24 for bisphenols) in different European states and harmonize measurements to make comparable data generated by different laboratories. HMB4EU will use these biomonitoring techniques to assess human exposure to chemicals in Europe and better understand health effects and improve chemical risk assessment. For the analysis of the metabolites of DEHP in the urine, there is no commercially available certified material to test an analytical method. Participation to the HBM4EU ICI/EQUAS tests allowed us to evaluate and validate analyses in urine samples, because provided a reference value from certified laboratories. The method here developed showed good sensitivity with LOQ ranging from 0.24 to 0.58 ng/ml, in line with methods previously published [83, 84, 86, 87]. For MEHP the lowest value provided was below the LOQ but the values obtained by our experimental method were very close to the reference values, in low and high concentrations. BIAS% was good ranging from 5.14 to 12.96 in the analyses conducted with different sample preparations, different times and different chromatographic runs, therefore including all possible variabilities. For BPA, on the other hand, we found numerous problems in LC/MS relating to the matrix effect in the analysis of urine samples; for this reason, we have developed a method that allows us to analyze BPA in GC/MS that allows good recovery and sensitivity. The LC/MS matrix effect affected the recovery of both analytes and internal standards and we observed a signal suppression likely due to the presence of molecules derived from enzymatic deconjugation that reduced the accuracy of the measurement. Indeed, the problem was evident in urine only after enzymatic deconjugation since the addition of different spikes of standard BPA did not result in a dose dependent recovery, while in free urine this matrix effect was not present. We showed that the matrix effect disappeared when the matrix analyzed by UHPLC/QTOF was derivatized and analyzed by GC/MS. In our results we demonstrated the importance of this work and that the majority of Italian children population (4–14 yrs old) and women were exposed to DEHP and BPA [97, 98], with measurable levels of its representative metabolites and showing a continuous and widespread exposure for DEHP (47.49 μg/L or 45.32 μg/g crea, as geometric mean, GM) as it emerged from LIFE PERSUADED project. Indeed, for DEHP metabolites both LC/MS and GC/MS methods have been reported, although analysis by LC/MS is the most common, while for BPA both instruments have been used although the accredited method (ISO17025) is by GC/MS/MS [99]. The HBM4EU proficiency test suggested to follow the method by GC/MS/MS validated in the LABERCA laboratory (France’s National Reference Laboratory for Food Residues and Contaminants) while no suggestions were given for phthalates. For both phthalates and BPA, we obtained good results within the acceptable ranges required by the ICI/EQUAS tests (Table 3). In conclusion the new method here presented has the advantage to start from low volumes of urine (0.5ml) and by using LC/QTOF followed by GC/MS allows the accurate measurement of both DEHP metabolites and BPA.

## 4. Materials and Methods

### 4.1 Assessment of urinary DEHP metabolites and BPA: the LIFE-PERSUADED method

We evaluated the methods previously published for the purification and quantification of phthalates and BPA in biological or non-biological matrix. Urinary BPA and DEHP metabolites are usually quantified by mass spectrometry, couple with either gas or liquid chromatography. The majority of the methods used LC/MS equipped with triple quadrupole for DEHP metabolites [83–89], but none at that time used a QTOF. For BPA both LC/MS and GC/MS were used. Our aim was to set up a LC/QTOF method to analyze the samples of the LIFE-PERSUADED project that aimed at the biomonitoring of phthalates (DEHP metabolites) and BPA in Italian healthy non obese children (age 4-14 years) and their mothers (900 couples) were collected during the human biomonitoring study [95, 97, 98]. Moreover, studies in idiopathic premature thelarche (IPT) and precocious puberty (IPP, a minimum of 30 girls each group, aged 2–7 years) and idiopathic obesity (IO, a minimum of 30 boys and 30 girls, aged 6–10 years) and in rats exposed to DEHP and BPA, at the same doses evaluated in the children of the project, to evaluate the effects of exposure in juvenile in vivo model. LC/QTOF high resolution mass spectrometry is the best method for discovery of known and unknown compounds in the matrix of interest; given its selectivity and accuracy it is becoming a useful tool to study the exposure to EDCs even in small biological sample [94]. Unlike LC/MS/MS system with target MRM acquisitions, the LC/QTOF system allows to acquire and quantify several compounds with high resolution in a single analysis compatibly with the chromatographic method. Since the LIFE-PERSUADED project enrolled children for whom (those 4-6 year old) it was sometime difficult to get a large sample size; thus, we developed a method that allowed to measure DEHP metabolites, BPA and creatinine concentrations starting from small sample size (0.5 ml for each analysis). Below we report the details and reliability of the method developed for the measurement of DEHP metabolites and BPA in urine samples starting from 0.5ml. More than 3000 samples were processed throughout the 4 year project including urine samples, standards and quality control showing good stability and reproducibility of the measurement over time (i.e., quality control samples run together with samples (Figure 4).

#### 4.1.1 Chemicals and preparation of standard solutions

Solutions and standard curves were prepared using unlabelled standards, i.e., Mono(2-ethylhexyl) phthalate (MEHP, ULM-4583-MT-1.2), Mono-2ethyl-5hydroxyhexyl phthalate (MEHHP, ULM-4662-MT-1.2) and mono-2ethy-5oxo hexyl phthalate (MEOHP, ULM-4663-MT-1.2) (Cambridge Isotope Laboratories, Tewksbury, MA); we used a nominal concentration of 100 ng/ml for each compound. As internal standards we used ^13^C-DEHP metabolites and prepared a mix solution containing 100 ng/ml of ^13^C4-MEHP (ring-1,2-^13^C2, dicarboxyl-^13^C2), 100 ng/ml ^13^C4-MEHHP and 100 ng/ml ^13^C4-MEOHP (Cam-bridge Isotope Laboratories, Tewksbury, MA). The standard solution of BPA (purchased from Dr. Ehrenstrofer Reference Materials Residue Analysis, Germany) was prepared by weighing and dissolving the compound in acetonitrile; the solutions concentration obtained was 535 ng/ml. d16BPA (C15 ^2^H16 O2, CDN isotopes, Pointe-Claire, Quebec CDN) was used as Internal Standard and was prepared by weighing and dissolving the compound in acetonitrile at final concentration of 216 ng/ml. The internal standards (IS) (40 μl of ^13^C DEHP metabolites and 20 μl of d16BPA, per sample) were added to the urine samples for quantification of compound concentrations.

#### 4.1.2. Preparation of standard curves

Standard solutions of MEHP, MEHHP, MEOHP were prepared in acetonitrile. For each metabolite standard curves were prepared 5 points with concentration of 4.1, 6.1, 8.1, 16.2 and 50.7 ng/ml of unlabelled compounds, that ideally covers the possible quantities pre-sent in the urine, and with 8 ng/ml of each internal standards. The final solution was dried under a gentle nitrogen flux and resuspended with 150 μl acetonitrile:water (1:9, v:v) as urine samples. Calibration curves were injected once a week before starting the analyses of urine samples and used to check linearity of the instrument in the defined concentrations and to correct the results for recovery. The standard curve of BPA was prepared using 7 different concentrations of BPA (from 1.1 to 71.5 ng/ml) and 20 μl of d16BPA (216 ng/ml) and was injected in GC/MS once a week before start sequences analysis.

#### 4.1.3 Preparation of Quality Control samples

For each metabolite we prepared and run Quality Control (Matrix QC) samples prepared by spiking the biological matrix (that was urine from a standard pool) with known amount of MEHP, MEOHP and MEHHP to reach a final concentration of 15.2 ng/ml for each metabolite and with BPA to reach 21.4 ng/ml. QC samples were prepared in batches before starting the analyses of the samples of the LIFE PERSUADED project and 200 μl aliquots were stored at −20°C for subsequent analyses. Repeated injections of matrix QC were made to evaluate method robustness and stability of frozen samples over a number of days.

#### 4.1.4. Collection and preparation of urine samples

Urine samples were collected into polypropylene (PP) tubes (i.e., without phthalate and BPA), centrifuged and stored at −20°C. Samples were thawed at 4 °C and 500 μl of urine was transferred in a glass tube. To each sample we added 375 μl of ammonium acetate buffer (NH4CH3COO-) 1M (Merck, Darmstadt, Germany), at PH=5, 250 μl of MilliQ water (Millipore Milli-Q Synthesis A10, Merck) and 2 μl of the enzyme β-glucuronidase (from Helix from Pomatia enzyme aqueous solution, ≥ 100.000 units/mL; Merck, Darmstadt, Germany) to deconjugate DEHP metabolites and BPA. The samples were incubated at 37°C overnight (for at least 16 hour). We also use Abalonase Purified β-Glucuronidase (United Chemical Technologies, Bristol, PA) >50,000 units/mL with 10x Rapid Hydrolysis Buffer (1:1, v:v with urine) and urine samples with mix labelled standard solution were incubated at 60°C for 1 hour. Instead after overnight incubation 25 μl of formic acid was added to stop the enzyme reaction and then 40 μl of the mix labelled standard solution was added to the samples. At last, 1 ml of MilliQ water was added into each sample to facilitate passage into solid phase extraction. For phthalates and BPA extraction we used Agilent Vac Elut Manifold with SPE cartridge C18 ODS 3 ml tubes 200 mg (Agilent Santa Clara CA, USA). SPE cartridges were conditioned with 2 ml of methanol and then 2 ml of MilliQ water. Urine samples were then loaded into the cartridges and washed with 1.5 ml of MilliQ water. Elution was performed with 1 ml of acetonitrile followed by 1 ml of methanol. The samples were then dried under a gentle N2 flux, reconstituted with 150 μl of acetonitrile:water (1:9, v:v) and transferred into glass vials for the analyses by UHPLC/QTOF [100]. We setup a GC/MS method that used the same de-glucuronidated urines previously prepared to measure DEHP metabolite concentrations by UHPLC/QTOF. Briefly, the samples were transferred to a glass tube and dried under a nitrogen flow. Then, 10 μl of BSTFA 1%TMS and 50 μl of acetonitrile (Merck, Darmstadt, Germany) were added to derivatize the samples and incubated at 75°C for 40 minutes. The sample was then transferred into glass vials avoiding contact with other materials for GC/MS analysis.

#### 4.2.1. Instrumental conditions

Analyses of DEHP metabolites by LC/QTOF. LC conditions: the LC system is an Ultra High Pressure (Agilent UHPLC 1290 infinity) with a binary pump, autosampler and Thermostatted Column Compartment (TCC). We used an Agilent ZORBAX SB-Phenyl column (2.1×100mm, 1.8-Micron, Agilent, Santa Clara CA, USA) maintained at 20°C with in-line Filter (0.3 μm SS Frit, 1.3 μl Delay Volume). The needle was washed with 100% methanol for 20 seconds and the injection volume was 5 μl. The mobile phase A was 10 mM ammonium acetate and the mobile phase B was methanol at a flow rate of 0.3 ml/min. The total run time was 14 minutes (including 2 minutes of equilibration time). The mobile phase gradient is reported in Table 4.

**Table 4.** Mobile phase gradient of the method: A= Water 10mM Ammonia Acetate; B=Methanol

| Time | Solvent A (%) | Solvent B (%) |
| --- | --- | --- |
| 0 | 67 | 33 |
| 4 | 67 | 33 |
| 6 | 40 | 60 |
| 7 | 5 | 95 |
| 10 | 67 | 33 |
| 12 | 67 | 33 |

##### MS conditions

Quadrupole Time-of-Flight (QTOF, Agilent 6540, Santa Clara CA) was equipped with the Dual ESI Jet Stream; the electrospray ionization was performed in negative mode. Instrument Mode was Extended Dynamic Range (2 GHz) in Mass Range Low (1700 m/z). The MS configuration and conditions are reported in details in Table 5. The acquisition mass range was m/z 100-500. Retention times were 7.6 min for MEHP, 4.4 min for MEOHP and 4.7 min for MEHHP

**Table 5.** QTOF parameters.

| Parameters | Values |
| --- | --- |
| Ionization mode | Negative ionization |
| Capillary voltage | -2,250 |
| Nozzle voltage | -300 |
| Nebulizer pressure | 35 |
| Dry gas temperature | 300 °C |
| Dry gas flow rate | 8 L/min |
| Sheath gas temperature | 350 °C |
| Sheath gas flow rate | 11 L/min |

Analyses of BPA by LC/ QTOF. Quantification of BPA in urines was first performed by UHPLC/QTOF simultaneously with phthalates using the conditions reported above. Although free BPA could be accurately quantified by LC/MS, for total BPA (i.e., after deconjugation) the analyses showed a matrix effect (described in the result session) that affected the reproducibility and accuracy of the measurements. Retention time for BPA was 6.8 minutes. Analyses of BPA by GC/MS. To assess total BPA, we used a GC/MS single quadrupole (GC 7890 - MS 5975, Agilent, Santa Clara, CA). Front SS Inlet was set in splitless mode at 280°C and carrier gas was Helium at constant flow rate of 2 ml/min. Oven program was 80 °C for 1 minute, then 30°C/min to 250°C for 0 minute and 40°C/min to 300°C for 2 minutes. Oven equilibration time was 0.5 minutes and run time was 9.9 minutes. Com-pounds were separated using a capillary column (DB-5MS J&W, l 30 m; i.d. 0.25 mm; film thickness 0.25 um). Samples were ionized by electron impact (EI) at 70 eV and chromatograms were acquired in selected ion monitoring (SIM). MS source was set up at 230°C and MS quad at 150°C and temperature in the transfer line was 280°C. The fragment ion for BPA was 357 m/z and for d16BPA was 368 m/z (Figure 6). Retention time for BPA was 6.8 min. and 6.78 for d16BPA in GC/MS.

**Figure 6.**
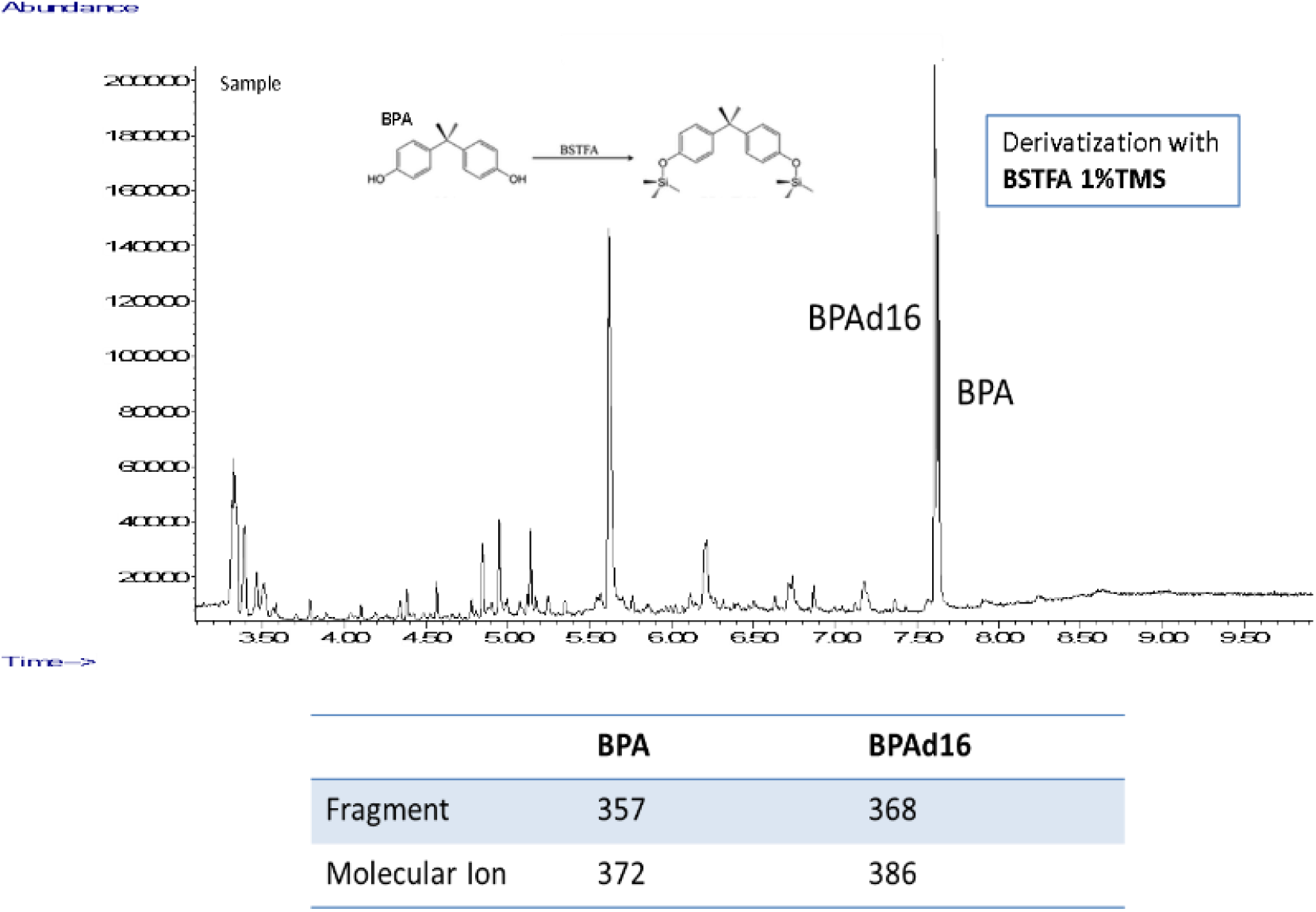
Chromatogram of BPA in GC/MS. BPA and BPAd16 were derivatized and the peak with 2 TMS groups was detected and quantified using fragment 357 m/z for BPA and 368 m/z for BPAd16 in SIM mode. Molecular ions were detected with fragment target ions for identification.

#### 4.2.2. Limits of Detection (LOD) and Quantification (LOQ)

Limits of Detection and Quantification (respectively LOD and LOQ) were established following the IUPAC guidelines ([101] also recommended by the U. S. Environment Protection Agency EPA (https://www.epa.gov/cwa-methods/procedures-detection-and-quantitation-documents, Office of Pesticide Programs U.S. Environmental Protection Agency Washington, DC 20460 March 23, 2000).

To calculate LOD and LOQ of phthalates we used the standard deviation of the first point of the curve that had a signal-to-noise ratio SNR>5 given by Agilent Mass Hunter Qualitative Analysis B.06.00 that calculates the distance from the height of the peak to the midline between the maximum and minimum noise at baseline. LOD and LOQ were set to 3 and 10 standard deviations above the response at 0 concentration (blank) estimated by the standard curve. For each metabolite (MEHP, MEHHP, MEOHP) we prepared 4 samples in acetonitrile:water (1:9, v:v) blank, concentration 0.1, 1, 3 and 5 ng/ml with a constant plus 8 ng/ml of ^13^C labelled standards. We also determined the Method Detection Limit (MDL), i.e., the minimum concentration of a substance that can be measured with a 99% confidence that the analyte concentration is greater than zero; MDL was calculated by multi-plying the standard deviation of lowest measurable sample by the Student’s t-value at the 99 percent confidence level.

LOD and LOQ for total BPA were calculated according to U.S.EPA procedure (U.S. EPA 2017). Briefly we assessed blank concentration of BPA during GC/MS runs by measuring standard deviation of more than 10 runs.

#### 4.2.3. Reagent blank and spike recovery

Reagent Blank and spike recovery were assessed by measuring DEHP metabolites and BPA in water and urine samples where known amount of labelled and unlabelled MEHP, MEOHP, MEHHP and BPA were added. Urines were extracted in SPE cartridge C18 (see preparation of urine samples). Recovery of DEHP metabolites was calculated as the ratio between the spike concentrations and nominal values (3.8 ng/ml). For LC/MS analysis matrix effect (ME) was calculated from ratio of concentration in spike urine sample and spike in water. For BPA analysis, before starting the sequence, the blanks with the internal standard (d16BPA) are run for about ten injections to settle the BPA contamination signal. Recovery of BPA was calculated as the ratio between concentration of spikes in urine and nominal values (10.4 ng/ml).

#### 4.2.4 Calculations of EDCs concentrations and assessment of exposure

Concentrations of DEHP metabolites and BPA were assessed with mass spectrometry and were normalized using creatine urine concentration to adjust the urinary concentrations of EDCs for the dilution. Results in LIFE PERSUADED project were reported un-adjust (μg/L) and adjusted (μg/g crea) to creatinine concentrations measured using the Jaffe method (Beckman Coulter, Brea, CA) [95, 98, 100, 102]. Assessment of DEHP exposure were estimated using sum of concentrations of its metabolites.

Relative metabolic rates (RMR) of the transformations from precursors to products (Figure 1) of MEHP, MEOHP and MEHHP were calculated as RMR1=([MEHHP]+[MEOHP])/[MEHP] and RMR2=([MEOHP]/[MEHHP]) x10, representing the rate of hydroxylation from MEHP to MEHHP and hydroxylation from MEHP to MEHHP respectively.

LC/MS chromatograms were analyzed with the instrumental software Mass Hunter Profinder B.06.00 (Agilent Technology, Santa Clara, CA) with targeted feature extraction (Figure 7) Mass Profinder algorithm was used to identify and extract compounds by known chemical formulas [M-H]- ±0.01 m/z tolerance since ionization was performed in negative mode (Figure 8), to calculate the relative isotope composition of compounds and peak area of each compound (Table 6). GC/MS chromatograms were analyzed by the GC/MSD ChemStation software (Agilent Technology, Santa Clara, CA).

**Figure 7.**
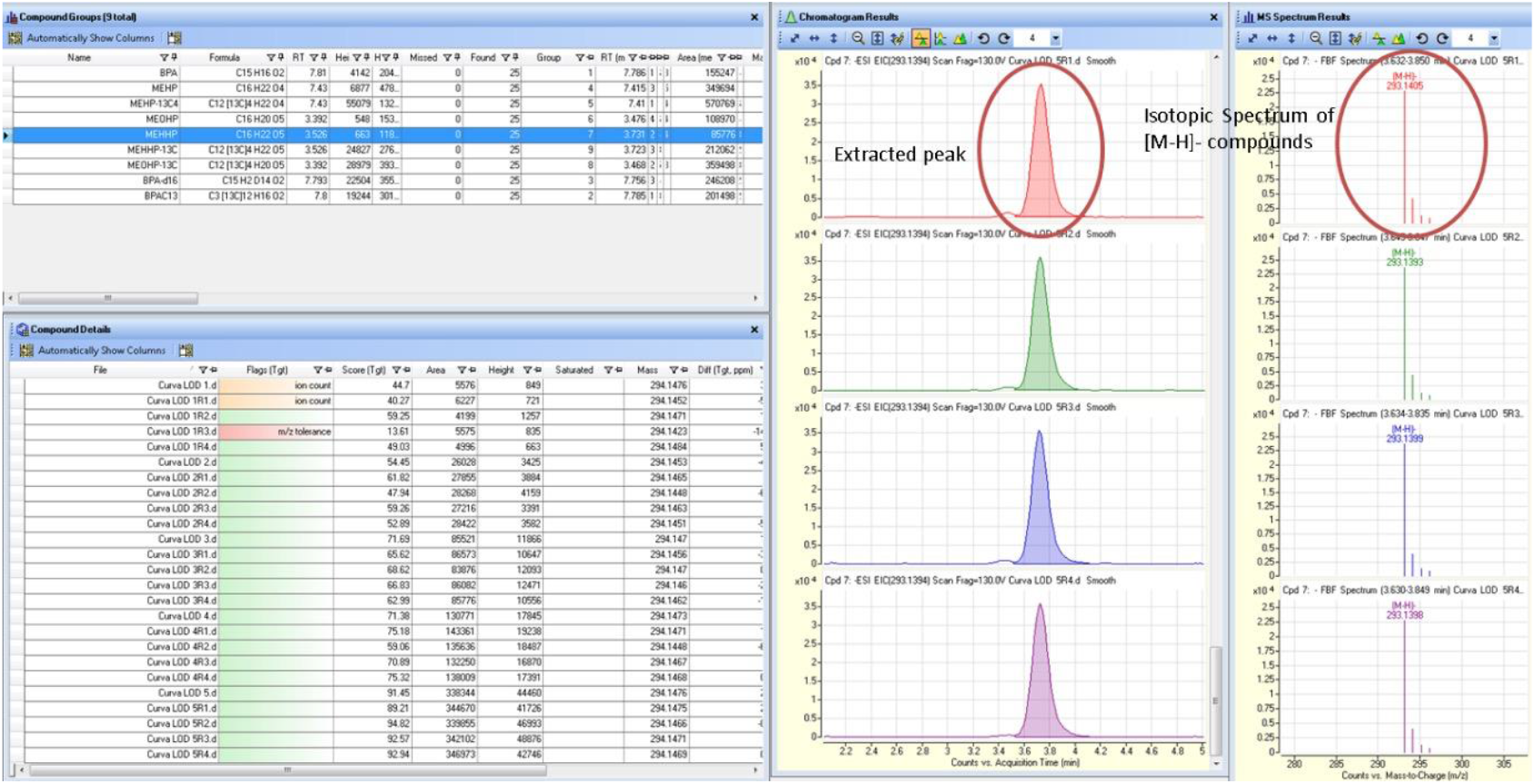
Chromatograms were analyzed with instrumental software Mass Hunter Profinder B.06.00 Profinder (Agilent, Santa Clara CA) with targeted feature extraction. Mass Profinder algorithm was used to identify and extract compounds by known chemical formulas ([M-H+] ±0.001 m/z tolerance

**Figure 8.**
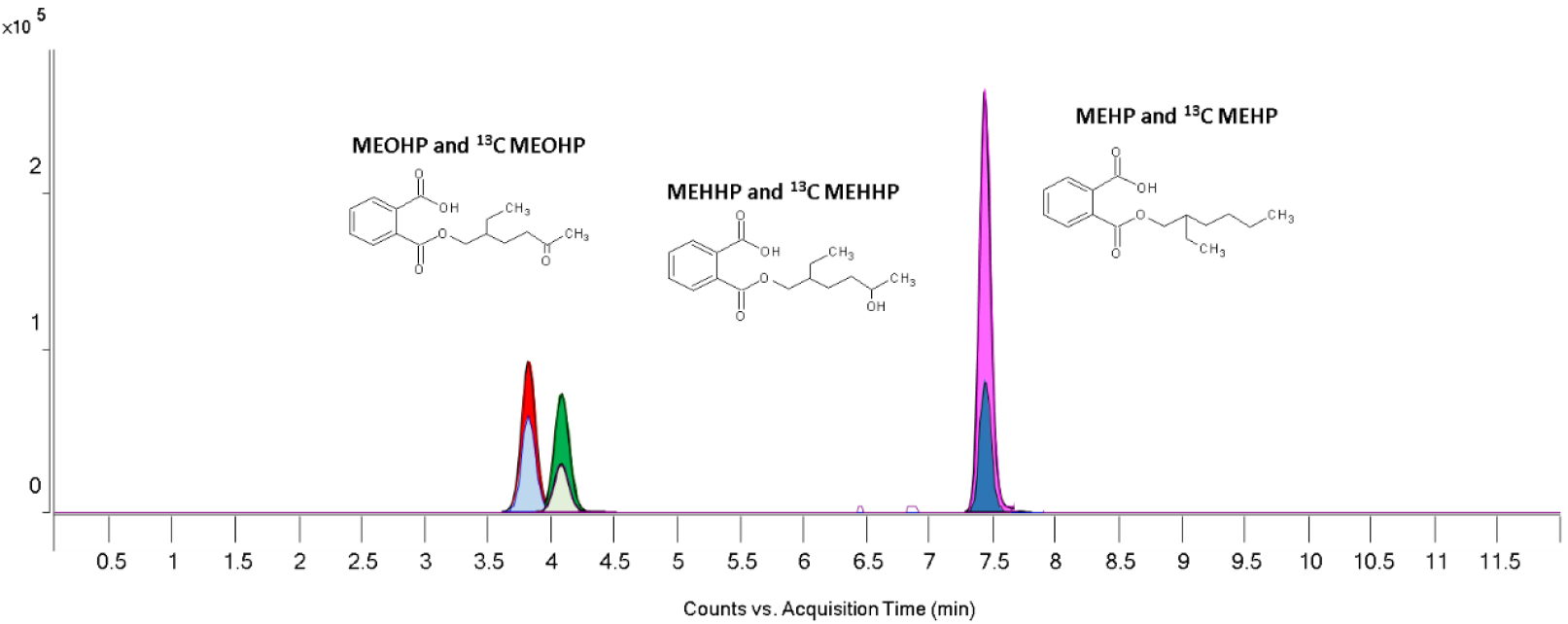
Peak extraction of phthalates metabolites and their ^13^C labelled standard by LC/ QTOF.

**Table 6.** Chemical features of MEHP, MEHOP, MEHHP and of their labelled standard compounds.

| Compound | Molecular Formula | MW | m/z $[M-H]^+$ |
| --- | --- | --- | --- |
| MEHP | $C_{16}H_{22}O_4$ | 278 | 277.1445 |
| $^{13}C_4$MEHP | $C_{12}[^{13}C]_4H_{22}O_4$ | 282 | 281.1580 |
| MEOHP | $C_{16}H_{20}O_5$ | 292 | 291.1238 |
| $^{13}C_4$ MEOHP | $C_{12}[^{13}C]_4H_{20}O_5$ | 296 | 295.1372 |
| MEHHP | $C_{16}H_{22}O_5$ | 294 | 293.1394 |
| $^{13}C_4$ MEHHP | $C_{12}[^{13}C]_4H_{22}O_5$ | 298 | 297.1529 |

The concentration of each compound was calculated from the peak area ratio of the labelled internal standards to the unlabelled compounds as below

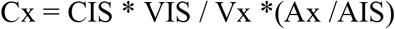

Where Cx = compound concentration; CIS = internal standard concentration; VIS=volume of IS; Vx=volume of compound x; Ax=compound area; AIS= Area of the internal standard compound.

## Acknowledgments

For drawing we used Servier Medical Art (smart.servier.com).

We acknowledge the LIFE PERSUADED Project consortium:

Istituto Superiore di Sanità, Rome, Italy: Cinzia La Rocca, Lucia Coppola, Gabriele Lori, Francesca Maranghi, Laura Narciso, Annalisa Silenzi, Sabrina Tait, Roberta Tassinari (Center for Gender-Specific Medicine); Luca Busani, Francesca Romana Mancini, Roberta Urciuoli (Department of Infectious Diseases, Istituto Superiore di Sanità); Antonio Di Virgilio, Andrea Martinelli, Mauro Valeri (Centro Nazionale Sperimentale Benessere Animale, Istituto Superiore di Sanità, Rome, Italy)

Dipartimento Pediatrico Universitario Ospedaliero “Bambino Gesu”, Rome, Italy: Stefano Cianfarani, Barbara Baldini Ferroli, Giorgia Bottaro, Annalisa Deodati, Romana Marini, Giuseppe Scirè, Gian Luigi Spadoni

Tor Vergata University, Rome, Italy; Stefano Cianfarani, Daniela Germani

Istituto di Fisiologia Clinica, CNR, Pisa, Italy: Amalia Gastaldelli, Graziano Barsotti, Emma Buzzigoli, Fabrizia Carli, Demetrio Ciociaro, Raffaele Conte, Veronica Della Latta, Graziella Distante, Melania Gaggini, Patrizia Landi, Anna Paola Pala, Chiara Saponaro.

Network of Italian pediatricians: https://lifp.iss.it/?p=73

